# Decoding the Centromeric Region with a Near Complete Genome Assembly of the Oshima Cherry *Cerasus speciosa*

**DOI:** 10.1101/2024.06.17.599445

**Authors:** Kazumichi Fujiwara, Atsushi Toyoda, Bhim B. Biswa, Takushi Kishida, Momi Tsuruta, Yasukazu Nakamura, Noriko Kimura, Shoko Kawamoto, Yutaka Sato, Toshio Katsuki, Sakura 100 Genome Consortium, Tsuyoshi Koide

## Abstract

The Oshima cherry (*Cerasus speciosa*), which is endemic to Japan, has significant cultural and horticultural value. In this study, we present a near complete telomere-to-telomere genome assembly for *C. speciosa*, derived from the old growth “Sakurakkabu” tree on Izu Oshima Island. Using Illumina short-read, PacBio long-read, and Hi-C sequencing, we constructed a 269.3 Mbp genome assembly with a contig N50 of 32.0 Mbp. We examined the distribution of repetitive sequences in the assembled genome and identified regions that appeared to be centromeric. Detailed structural analysis of these putative centromeric regions revealed that the centromeric regions of *C. speciosa* comprised repetitive sequences with monomer lengths of 166 or 167 bp. Comparative genomic analysis with *Prunus sensu lato* genome revealed structural variations and conserved syntenic regions. This high-quality reference genome provides a crucial tool for studying the genetic diversity and evolutionary history of *Cerasus* species, facilitating advancements in horticultural research and the preservation of this iconic species.

## Background & Summary

Oshima cherry [*Cerasus speciosa* (Koidz.) H.Ohba and *Prunus speciosa* (Koidz.) Nakai] is a cherry tree species endemic to Japan, found primarily in the wild on the Izu Islands ^1^. The classification of flowering cherry regarding the nomenclature of the genera as *Prunus* or *Cerasus* remains a controversy. However, we discuss the Oshima cherry and other flowering cherries within the genus *Cerasus* based on recent molecular and population genetic analyses ^2^. Flowering cherries have held significant cultural importance in Japan for millennia, not only utilized as a source of firewood, timber for buildings, and material for cords, but also cherished for their high ornamental value since the late 8th century ^3-5^. This appreciation has continued to the contemporary “Hanami” culture, a tradition of cherry blossom viewing that persists in Japan. Japan is native to ten species of wild cherry trees ^1,6-8^, and the number of named cultivars exceeds several hundred ^9-11^. Numerous cultivars within the “*Cerasus* Sato-zakura Group” are presumed to be descendants of wild Oshima cherry ^12^. Additionally, *Cerasus* ×*yedoensis* ‘Somei-yoshino’, one of the most famous Japanese cherry cultivars and an interspecific hybrid of *C. itosakura* and *C. speciosa*, is speculated to have one haplotype derived from the wild Oshima cherry lineage ^13-15^. The Oshima cherry stands out among Japan’s native wild species because of its diverse blooming periods, relatively large petal size, rapid growth, and strong fragrance derived from coumarins ^16^. These characteristics have made the Oshima cherry highly favored, significantly influencing the development of cherry cultivars in Japan ^17^. Both wild and cultivated Japanese cherry species are primarily diploids (2n = 2x = 16), possessing eight pairs of chromosomes similar to the karyotype of *C. speciosa* ^18^. Analysis of the Oshima cherry genome may greatly aid in understanding the genetics of the Japanese cherry cultivar.

Izu Oshima, one of the Izu Islands, houses a giant Oshima cherry tree, referred to as “Sakurakkabu,” which is estimated to be over 800 years old ^19^. This particular *C. speciosa* tree has been protected as a Natural Monument of Japan since 1935. This tree was originally a single massive tree until the Edo period (1603–1868), but its main trunk eventually broke. While multiple sprout trunks emerged from the remaining main trunk, they were repeatedly toppled due to the impact of numerous typhoons. From its remains, branches of various sizes spread out in all directions, with some growing diagonally to the ground or lying horizontally and rooting from points of contact, before growing upward again, forming a unique tree shape. As a result, the tree now comprises the original trunk and three additional trunks, each pointing north, east, and west. “Sakurakkabu” is believed to be the oldest existing Oshima cherry tree in Izu, Japan. Decoding the genome of this Oshima cherry tree may contribute significantly to the research on the origin of Oshima cherry and Japanese cherry trees. Furthermore, our findings may benefit studies of cultivars in Japan and other regions.

Herein, we report the near complete genome assembly of the Oshima cherry, *C. speciosa.* Samples of *C. speciosa* were obtained from the leaf buds of the original trunk of “Sakurakkabu” on Izu Oshima, with the permission of the Tokyo Metropolitan Government and the Agency for Cultural Affairs. By extracting genomic DNA and sequencing it using Illumina short reads and PacBio long reads, we constructed a high-quality telomere-to-telomere reference genome for *C. speciosa*.

## Methods

### Sample collection and sequencing

Young leaves collected from the Oshima cherry tree “Sakurakkabu” in Izu Oshima, Tokyo, Japan, were used for genomic DNA extraction (Fig. 1a). The leaves were collected, rinsed with deionized water, blotted dry with paper towels, and stored at [80 °C until the extraction of genomic DNA. Extraction of genomic DNA from the leaves was performed with the method reported previously with some modifications ^20^. Briefly, 0.1–0.5 g of leaves were ground into a powder in liquid nitrogen using a chilled pestle and mortar. This powder was suspended in 10 mL of Carlson lysis buffer (100 mM Tris-HCl pH 9.5, 2% CTAB, 1.4 M NaCl, 1% PEG 6000, 20 mM EDTA) supplemented with 0.25% β-mercaptoethanol, and then incubated at 74 °C for 20 min with gentle mixing by inverting the tube slowly every 5 min. The mixture was cooled to room temperature, followed by the addition of 20 mL chloroform/isoamyl alcohol (24:1), careful and gentle mixing, and centrifugation at 1,800 G for 10 min. The supernatant was transferred to a new tube, and an equal volume of 2-propanol were added. The mixture was mixed uniformly and centrifuged at 2,500 G for 30 min. The supernatant was discarded, and the cells were washed with 75% EtOH, followed by centrifugation for 10 min. The supernatant was discarded again, and the cells were briefly air-dried on paper towels. Subsequently, 1 mL of TE was added to dissolve the pellet. Then RNase was added to 10∼100 µg/mL and the sample was stored overnight at 4 °C. Genomic DNA was purified using a QIAGEN Genomic-tip 500G and QIAGEN DNA Buffer Set (QIAGEN, Hilden, Germany), following the manufacturer’s protocols. For sequencing using the Illumina NovaSeq 6000 platform with 250 bp paired-end reads, the genomic DNA was fragmented using Covaris, followed by library preparation using the TruSeq DNA PCR-Free Library Prep Kit. Size selection was performed via agarose gel excision. Sequencing was performed for 500 cycles using a NovaSeq 6000 Reagent Kit. For the sequencing of continuous long reads (CLR) using the Pacific Biosciences (PacBio) Sequel II platform, the genomic DNA was fragmented using g-TUBE. Library preparation was performed using the SMRTbell Express Template Prep Kit 2.0, followed by the removal of DNA fragments under 40 kb using BluePippin. Sequencing was performed using Sequel II Binding Kit 2.0/Sequencing Kit 2.0. For the sequencing of high-fidelity (HiFi) reads using the Pacific Biosciences (PacBio) Sequel II platform, the genomic DNA was sheared into fragments of 10–30 kb using a g-tube device (Covaris Inc., MA, USA). A SMRTbell library for HiFi reads was prepared using the SMRTbell Prep Kit 3.0 (Pacific Bioscience, CA, USA), according to the manufacturer’s instructions. The HiFi library was size-selected for 15-30 kb DNA fragments using the Sage ELF system (Saga Science, MA, USA) and sequenced on one SMRT Cell (8M) using the Sequel II system with the Binding Kit 3.2/Sequencing Kit 2.0 (Pacific Bioscience, CA, USA), with 1800 min collection times. Consensus HiFi reads were generated from raw full-pass subreads using the DeepConsensus v1.2 program ^21^. The Hi-C library was prepared using the Dovetail Omni-C Kit (Dovetail Genomics, CA, USA), according to the manufacturer’s protocol. Briefly, the chromatin was fixed in nucleus by disuccinimidyl glutarate (DSG) and formaldehyde. The crosslinked chromatin was digested using DNase I, and the cells were lysed to extract chromatin DNA fragments. The chromatin DNA ends were repaired and ligated to a biotinylated bridge adapter, and then proximity ligated of the adapter-containing ends. Subsequently, the cross-linked DNA was reversed and converted into a sequencing library using Illumina-compatible adaptors. Biotin-containing fragments were enriched with streptavidin magnetic beads before PCR amplification. The concentration and quality of the libraries were evaluated using a Qubit 4 fluorometer (Thermo Fisher Scientific, MA, USA), 2100 Bioanalyzer System (Agilent Technologies, CA, USA), and 7900HT Fast Real-Time PCR System (Thermo Fisher Scientific, MA, USA). Finally, the Omni-C library was sequenced on an Illumina NovaSeq 6000 system with a 2 × 150 bp read length, generating 61.2[Gb of Omni-C reads.

**Figure 1.**
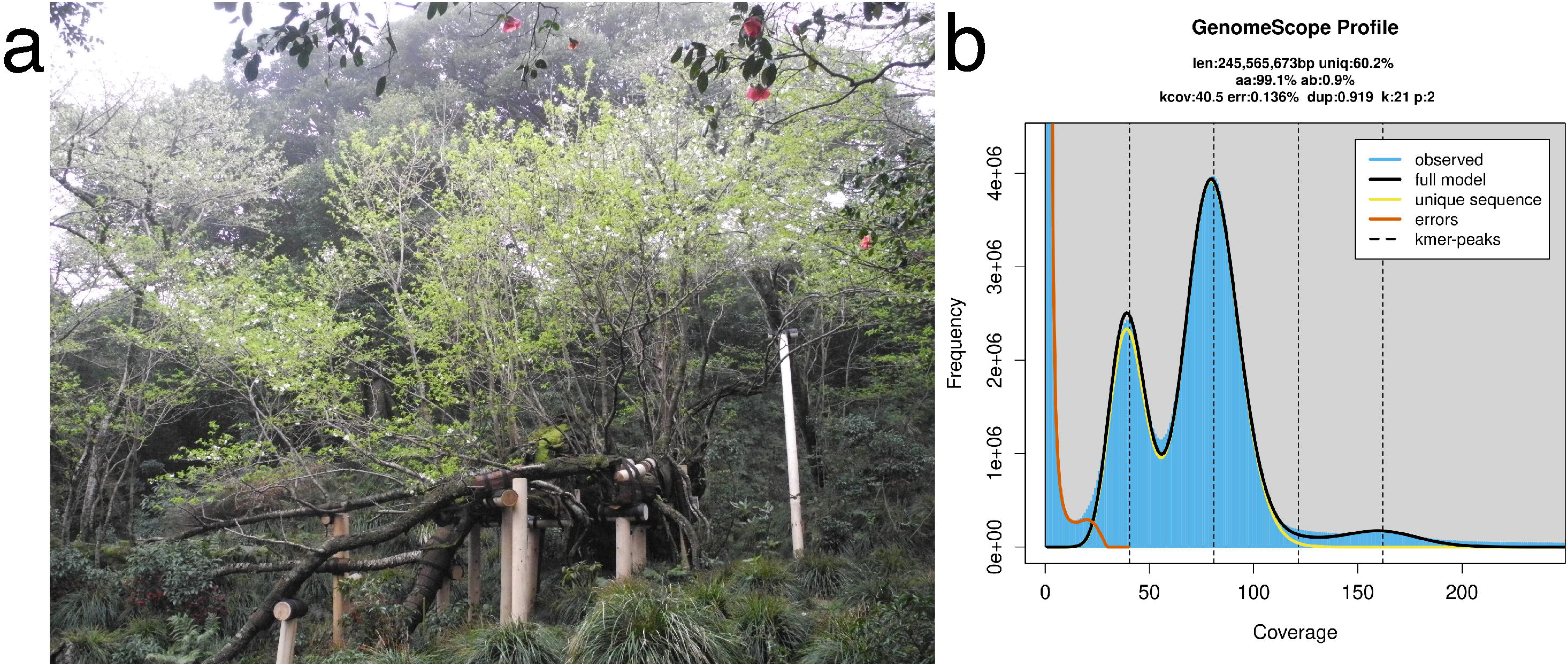
(a) Oshima cherry tree “Sakurakkabu” of Oshima, *Cerasus speciosa*. (b) Genome size estimation and heterozygosity through k-mer analysis.

RNA was extracted separately from the homogenized leaf buds, floral buds, stamens, and ovules of *C. speciosa*, which were processed in liquid nitrogen. The resulting frozen powder (10–20 mg) was used for RNA extraction, employing the Fruit-mate™ and NucleoSpin™ RNA Plant kits (Takara Bio Inc., Shiga, Japan), in accordance with the manufacturers’ instructions. RNA quality and concentration were assessed using an Agilent 2100 Bioanalyzer (Agilent Technologies, CA, USA) with the RNA6000 Pico kit (Agilent Technologies, CA, USA). Subsequently, the RNA samples were reverse-transcribed to cDNA using the SMART-Seq HT Kit (Takara Bio USA, CA, USA) following the manufacturer’s guidelines. Sequencing libraries were generated from the cDNA using the Nextera XT DNA Library Preparation Kit (Illumina, CA, USA) and sequencing was conducted on an Illumina NovaSeq 6000 system with a 2 × 100 bp read length.

### Genome size and heterozygosity estimation

The genome size of *C. speciosa* was estimated using *k*-mer based method and Illumina short-read sequencing datasets. The acquired short reads were subjected to rigorous quality control using fastp v0.23.2 ^22^ with the parameter “-q 30 -f 5 -F 5 -t 5 -T 5 -n 0 -u 20.” To manipulate and count the *k*-mer in short reads dataset, we utilized KMC3 v3.2.1 ^23^ counting canonical 21-mers with the parameter “-k 21 -ci 1 -cs 10000.” Our decision to select *k* = 21 was based on recommendations for genome size estimation processes, where *k* = 21 is advocated. This recommendation is based on the premise that a *k*-mer length of *k =* 21 is sufficiently long to ensure that most *k*-mer are not repetitive, yet short enough to enhance the robustness of the analyses against sequencing errors. For generating a histogram of *k*-mer statistics, kmc_tools implemented in KMC3 were executed by operation “transform” and “histogram” with parameter “-cx 10000.” The obtained histogram was analyzed by the Rscript version of GenomeScope 2.0 v2.0 ^24,25^ to estimate the haploid genome size and heterozygosity from *k-*mer count dataset with parameter “-k 21 -p 2.” The majority of cherry blossoms in Japan are diploid, and karyomorphological research has also confirmed that *C. speciosa* is diploid ^18^. Consequently, a diploid model (-p 2) was employed for genome size estimation using GenomeScope 2.0. Based on *k*-mer analysis, the haploid genome size of *C. speciosa* was estimated to be ∼245.6 Mb with a heterozygosity of 0.91% and a repeat content of 39.8% (Fig. 1b).

### De novo genome assembly and quality assessments

Primary genome assembly was performed using Hifiasm v0.19.5-r587 ^26,27^, incorporating PacBio HiFi reads along with Omni-C Illumina short reads. HiFi reads were generated from subread output by the Sequel II system using DeepConsensus v1.2 ^21^, with default parameters and then assembled with Omni-C data using Hifiasm under default settings. The number of HiFi reads was adjusted to optimize the assembly, ultimately using 85% for improved contiguity. Duplicate regions in the assembly were identified and removed using purge-dups ^28^, with the mapping process facilitated by minimap2 ^29,30^. Contamination was removed by filtering out contigs with more than 10% of the length aligning to the NCBI Prokaryotic RefSeq Genomes or the organelle genome of *C. speciosa* using blastn ^31^, with an e-value threshold of “1e-10.”

Finally, we obtained a complete genome assembly spanning 269,259,082 bp, characterized by a contig N50 value of 32.0 Mbp and GC ratio of 38.1%, with no gaps within the eight pseudomolecules corresponding to the chromosomes (Fig.2, Table 1).

**Figure 2.**
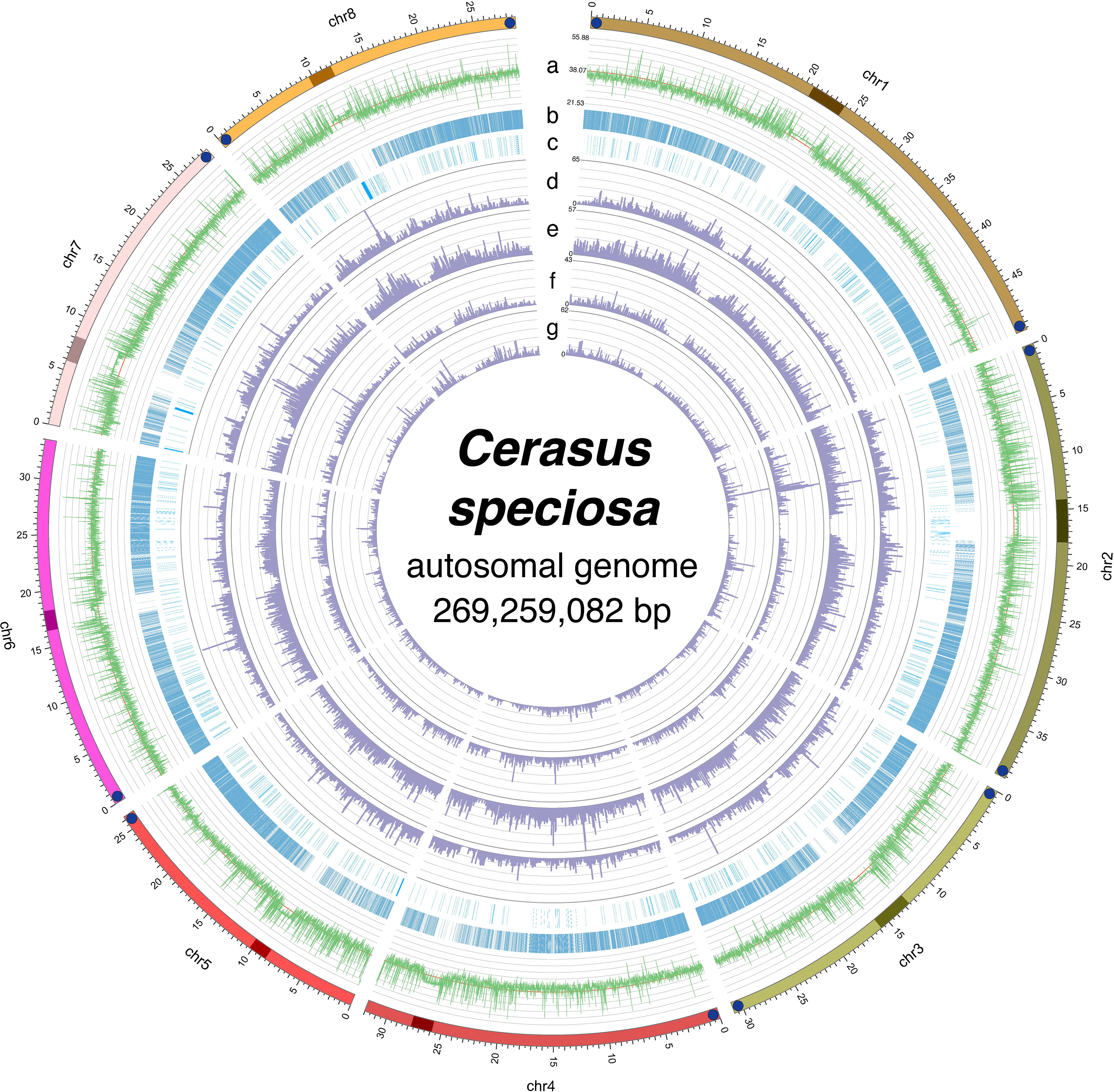
Schematic illustration of the genetic features of the *C. speciosa* genome. The outermost circle represents the length of each chromosome, with telomere sequences at the ends (blue circles) and estimated centromeric regions (darker color). From the outermost to the innermost track, each circle represents the following genetic characteristics: guanine-cytosine (GC) ratio (a), gene distribution (b), non-coding RNA distribution (c), Long Terminal Repeats (LTR) (d), Terminal Inverted Repeats (TIR) (e), non-LTR elements (LINE and SINE) (f), and non-TIR elements (Helitron) (g).

**Table 1.**
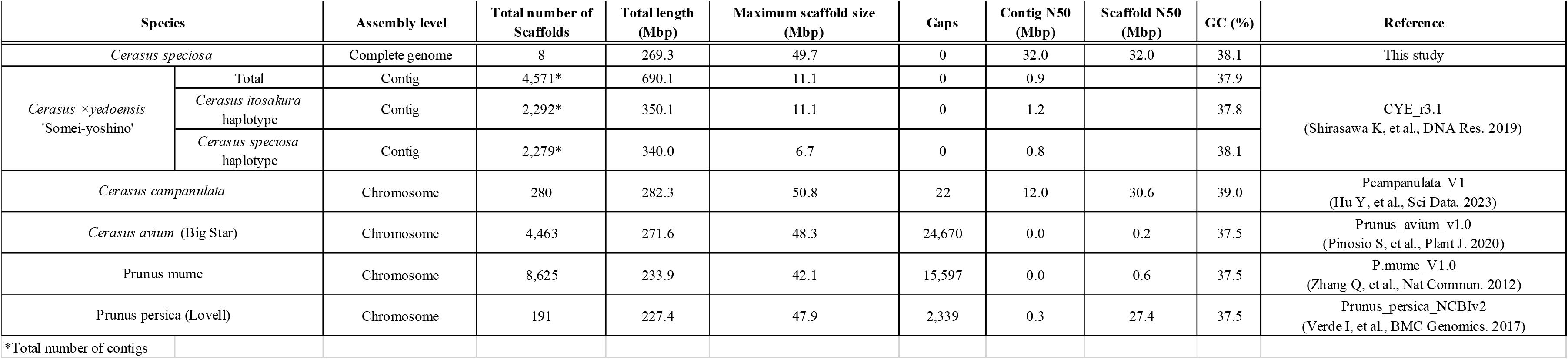
Summary statistics of the *Cerasus speciosa* genome assembly and other closely related species genomes.

To evaluate our newly assembled genome of *C. speciosa*, we used Benchmarking Universal Single-Copy Orthologs (BUSCO) v5.5.0 ^32^ to assess the completeness of the genome. The BUSCO lineage dataset was downloaded from OrthoDB v10 ^33,34^. To determine the dataset most suitable for the *C. speciosa* genome, the “--auto-lineage” option was specified, using the eukaryota_odb10 and eudicots_odb10. The BUSCO assessments revealed completeness of 99.6% (254 out of 255 genes) for the eukaryota dataset and 98.4% (2289 genes out of 2326 genes) for the eudicots dataset (Table 2). The BUSCO datasets were evaluated further using compleasm v0.2.4 ^35^ by “run” mode with “-l eudicots” option. Compleasm is recognized as a more rapid and accurate tool for reassessment than BUSCO. Using the same eudicots dataset as in the previous BUSCO analysis for evaluating genome assemblies, we observed a completeness of 99.14% (2,306 of 2,326 genes), with a composition of 97.03% single-copy complete genes (2,257 genes), 2.11% duplicated complete genes (49 genes), 0.39% fragmented genes (9 genes), and 0.47% missing genes (11 genes) (Table 2). In addition to the conventional evaluation of genome assemblies using BUSCO, we utilized the methodology known as Inspector v1.0.1 ^36^, which assesses genome assemblies using long reads. This approach involves aligning the long reads used for genome assembly with the assembled genome to evaluate the completeness of the genome structure. In this study, we conducted evaluations using PacBio HiFi reads (“-d hifi” option) employed in the assembly process. The mapping rate of the long reads to the assembled genome was 99.99%, with a quality value (QV) of 35.89, indicating an accuracy of 99.97% (Table 2). Furthermore, we used a *k*-mer-based approach to evaluate genome assembly with Merqury v1.3 (Rhie et al., 2020). For the *k*-mer counting, Meryl v1.3, implemented in Merqury, was used to compute the *k*-mers from PacBio HiFi reads with *k =* 21 settings. Merqury was proceeded with its default parameters for pseudo-haplotypes using the *k*-mer datasets quantified by Meryl. The results indicated a QV > 67, corresponding to an error rate of 1.80 × 10^-7^ (Table 2).

**Table 2.** Genome assembly and annotation statistics for *C. speciosa*.

| **Genome assembly statistics** | **Value** |
| --- | --- |
| BUSCO analysis |  |
| lineage dataset | eudicots_odb10 |
| Completeness | 98.4% (2289/2326) |
| Complete single-copy | 95.9% (2231/2326) |
| Complete duplicated | 2.5% (58/2326) |
| Fragmented | 0.5% (11/2326) |
| Missing | 1.1% (26/2326) |
| Compleasm analysis |  |
| lineage dataset | eudicots_odb10 |
| Completeness | 99.1% (2306/2326) |
| Complete single-copy | 97.0% (2257/2326) |
| Complete duplicated | 2.1% (49/2326) |
| Fragmented | 0.4% (9/2326) |
| Missing | 0.5% (11/2326) |
| Inspector analysis |  |
| reads-to-contigs mapping rate | 99.99% |
| QV | 35.89 (99.97% accuracy) |
| Merqury analysis |  |
| QV | QV>67 |
| error rate | $1.80 \times 10^{-7}$ |

### Assembled genome annotations

To annotate repetitive sequences, we employed EDTA v2.0.0 ^37^ to conduct annotation using *de novo* and curated libraries using a homology-based approach. For the curated libraries, we employed a collection known as nrTEplants v0.3 ^38^, which contains repetitive sequences annotated from various databases, including REdat ^39^, RepetDB ^40^, TREP ^41^, and other collections. In conducting the EDTA analysis, coding sequences (CDS) from closely related species were provided, using the CDS of the *C. speciosa* haplotype from the already published genome of *C.* ×*yedoensis* ‘Somei-yoshino’ ^15^. The parameters employed in the execution of EDTA were as follows: --genome [*C. speciosa* genome] --cds [*C. speciosa* haplotype of *C.* ×*yedoensis* ‘Somei-yoshino’] --curatedlib [nrTEplants curated library] --overwrite 1 --sensitive 1 --anno 1 --evaluate 1. Through the annotation of repetitive sequences, approximately 128.8 Mb of repetitive regions were identified, accounting for 47.8% of the *C. speciosa* genome (Fig. 2, Table 3). Each type of repetitive sequence was classified using RepeatMasker v4.1.1 (https://www.repeatmasker.org/), which was implemented in an EDTA Docker image (oushujun/edta:2.0.0).

**Table 3.**
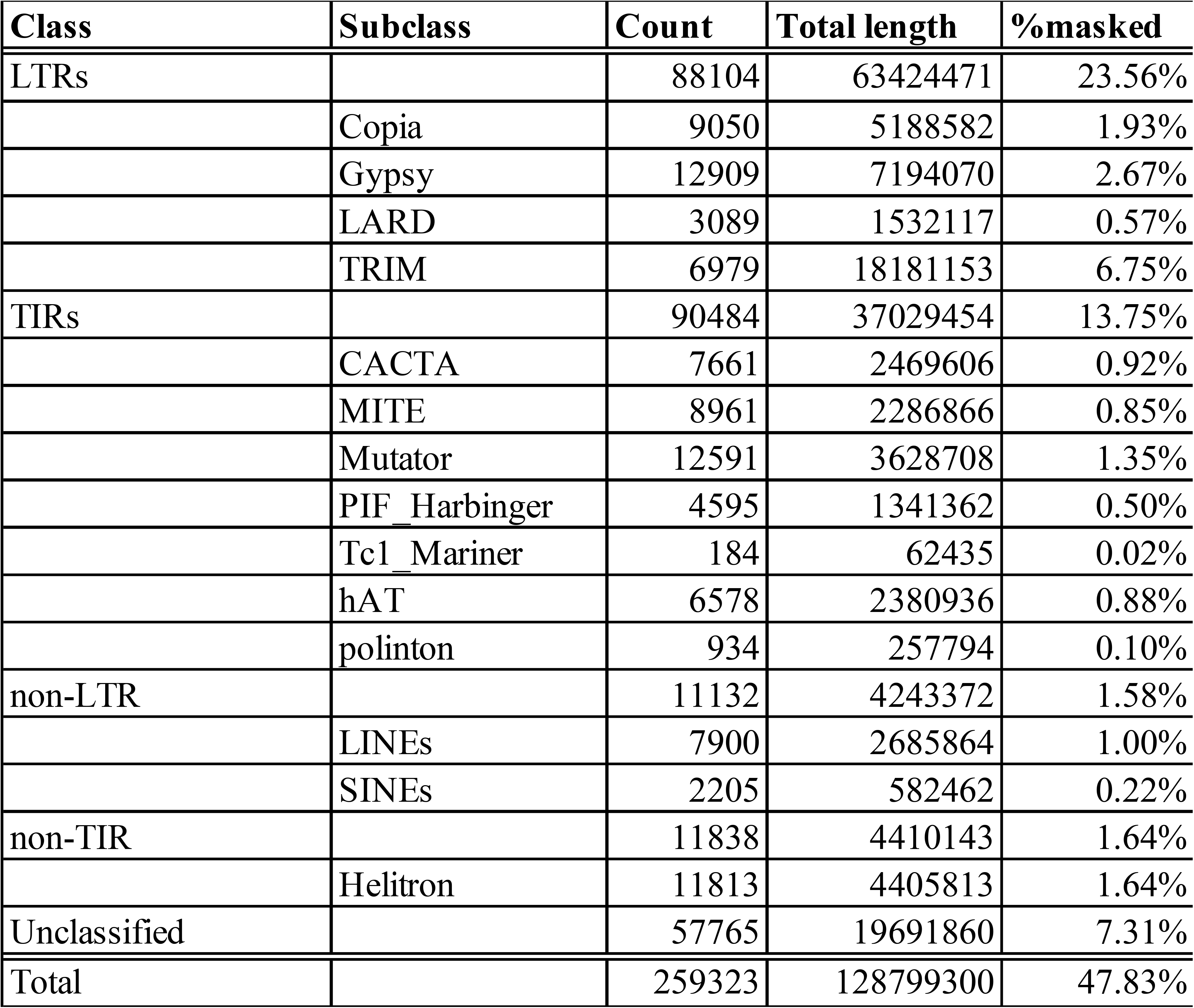
Summary of repetitive contents in *C. speciosa*.

| **Class** | **Subclass** | **Count** | **Total length** | **%masked** |
| --- | --- | --- | --- | --- |
| LTRs |  | 88104 | 63424471 | 23.56% |
|  | Copia | 9050 | 5188582 | 1.93% |
|  | Gypsy | 12909 | 7194070 | 2.67% |
|  | LARD | 3089 | 1532117 | 0.57% |
|  | TRIM | 6979 | 18181153 | 6.75% |
| TIRs |  | 90484 | 37029454 | 13.75% |
|  | CACTA | 7661 | 2469606 | 0.92% |
|  | MITE | 8961 | 2286866 | 0.85% |
|  | Mutator | 12591 | 3628708 | 1.35% |
|  | PIF_Harbinger | 4595 | 1341362 | 0.50% |
|  | Tc1_Mariner | 184 | 62435 | 0.02% |
|  | hAT | 6578 | 2380936 | 0.88% |
|  | polinton | 934 | 257794 | 0.10% |
| non-LTR |  | 11132 | 4243372 | 1.58% |
|  | LINEs | 7900 | 2685864 | 1.00% |
|  | SINEs | 2205 | 582462 | 0.22% |
| non-TIR |  | 11838 | 4410143 | 1.64% |
|  | Helitron | 11813 | 4405813 | 1.64% |
| Unclassified |  | 57765 | 19691860 | 7.31% |
| Total |  | 259323 | 128799300 | 47.83% |

The non-coding RNAs (ncRNA) were annotated using three primary sources, tRNAscan-SE 2.0 v2.0.12 ^42^ for transfer RNA (tRNA); RNAmmer v1.2 ^43^ for ribosomal RNA (rRNA); and Rfam 14.10 ^44^ for ncRNAs, including microRNAs (miRNAs) from miRbase ^45^. We detected tRNA using tRNAscan-SE 2.0 with default options. We used RNAmmer to detect rRNA sequences with the following options specified: “-S euk -m lsu,ssu,tsu -multi.” miRNAs and other ncRNAs were identified using Infernal v1.1.4, which was predicted based on the Rfam database. The parameters for Infernal were specified in accordance with the guidelines documented in Rfam 14.10, and the final E-value cutoff for ncRNA was set to “1e-10.” Our predictions identified 572 tRNA genes encoding 20 amino acids, 24 tRNA pseudogenes, 794 rRNA genes, and 623 other ncRNAs (Fig. 2).

While annotating protein-coding genes, the initial step involved soft-masking repetitive sequences within the genome using RepeatModeler2 v2.0.5 ^46^ and RepeatMasker v4.1.5, implemented in the Dfam Transposable Element Tools Docker container v1.87 (https://github.com/Dfam-consortium/TETools). We constructed the repeat library using RepeatModeler2 incorporating the “-LTRStruct” option to engage the LTR discovery pipeline. For the soft-masking genome sequence using RepeatMasker, the following parameters were employed: “-pa 12 -a -html -gff -s -no_is -norna -nolow -div 40.” For the structural annotation of protein-coding genes, the BRAKER3 v3.0.7.1 ^47,48^ pipeline was employed, integrating predictions from three strategies: *ab initio* prediction, transcription evidence from RNA-seq, and homologous proteins from closely related species. To maximize the gene annotation predictions using RNA-seq, we implemented a noise-reduction process on the RNA-seq data. Specifically, the quality of the RNA-seq data was assessed using fastp with the following parameters: “-q 30 -f 5 -F 5 -t 5 -T 5 -n 0 -u 20.” We performed *de novo* transcriptome assembly using Trinity v2.15.1 ^49^ and mapped the RNA-seq data to this assembled genome using Hisat2 v2.2.1 ^50^. Only reads that were correctly mapped to the *de novo* assembled transcriptome genome were used for downstream analysis. During this process, we selected the genome-guided assembly option in Trinity, with a maximum intron size of 1000 bp. The rationale for setting the maximum intron size to 1000 bp was based on the average and median intron sizes of less than 1000 bp in the genomes of closely related species (*Prunus persica, Prunus mume,* and *Cerasus avium* (formerly named *Prunus avium*)) registered in the NCBI RefSeq database. Additionally, the “--very-sensitive” option was utilized in Hisat2 to enhance mapping sensitivity. The extracted reads were then re-mapped to the complete *C. speciosa* genome using Hisat2, which served as an input file for BRAKER3. For this remapping, Hisat2 was configured with the “--very-sensitive” and “--max-intronlen 15000” options. For the structural annotation of protein-coding genes using BRAKER3, the “--AUGUSTUS_ab_initio” option was used to perform *ab initio* gene predictions using AUGUSTUS ^51^, implemented in BRAKER3. Furthermore, protein sequences from the *C. speciosa* haplotype of *C.* ×*yedoensis* ‘Somei-yoshino’ were provided as the required protein information from a closely related species for BRAKER3. The RNA-seq data used as input files for BRAKER3 were obtained using previously described processes. To enhance the annotation results from BRAKER3, the “--busco_lineage=eudicots_odb10” option was selected for BUSCO assessment, utilizing Compleasm ^35^ implemented in BRAKER3. The annotation of protein-coding genes by BRAKER3 does not include information on untranslated regions (UTRs). Therefore, to supplement this deficiency, GUSHR v1.0.0 (https://github.com/Gaius-Augustus/GUSHR) was executed using the RNA-seq data. To predict the functions of genes inferred by structural annotation using BRAKER3, functional annotation was performed using EnTAP v1.0.1 ^52^. EnTAP performed homologous alignments using DIAMOND v2.1.8.162 ^53^ across multiple databases, including SwissProt, TrEMBL, and the non-redundant plant protein database of NCBI (NR). By predicting the structural and functional annotations of protein-coding genes, we predicted 33,773 protein-coding genes from the *C. speciosa* genome, of which 29,776 genes were functionally annotated against public databases (Fig. 2). The BUSCO completeness of the protein sequences was 97.4% (n = 2267) in “eudicots_odb10” lineage (n = 2326), including 78.3% (n = 1822) single-copy, 19.1% (n = 445) duplicated, 0.3% (n = 8) fragmented, and 2.3% (n = 51) missing BUSCOs. The compleasm completeness of the protein sequence using BUSCO gene set was 97.5% (n = 2268) in “eudicots_odb10” lineage (n = 2326), including 69.7% (n = 1622) single-copy, 27.8% (n = 646) duplicated, 0.7% (n = 16) fragmented, and 1.8% (n = 42) missing BUSCOs. In both evaluations, the annotations were of very high quality.

### Chloroplast genome assembly and annotation

The chloroplast genome of *C. speciosa* was *de novo* assembled using Illumina short reads. Short reads were filtered and trimmed using fastp, as previously mentioned. Clean reads were then assembled using the get_organelle_from_reads.py script implemented in GetOrganelle v1.7.7.0 ^54^, with the parameters set to “-R 15 -k 21,55,85,115,127 -F embplant_pt.” Among the assemblies, one with a *k*-mer size of *k*=127 was selected, resulting in a complete circular chloroplast genome sequence of 157,895 bp in length. Gene annotation of the assembled complete chloroplast genome was performed using GeSeq (https://chlorobox.mpimp-golm.mpg.de/geseq.html) ^55^. For the configuration of GeSeq, the default settings were primarily utilized. However, modifications were made to the following options: the “FASTA file to annotate” was set to circular, the “sequence source” was changed to “plastid (land plants),” and *C. speciosa* (NCBI RefSeq: NC_043921.1) was employed as the reference sequence for BLAT analysis. Furthermore, the results of gene annotation were visualized as vector files using the OGDRAW web service (https://chlorobox.mpimp-golm.mpg.de/OGDraw.html) ^56^ (Fig. 3a).

**Figure 3.**
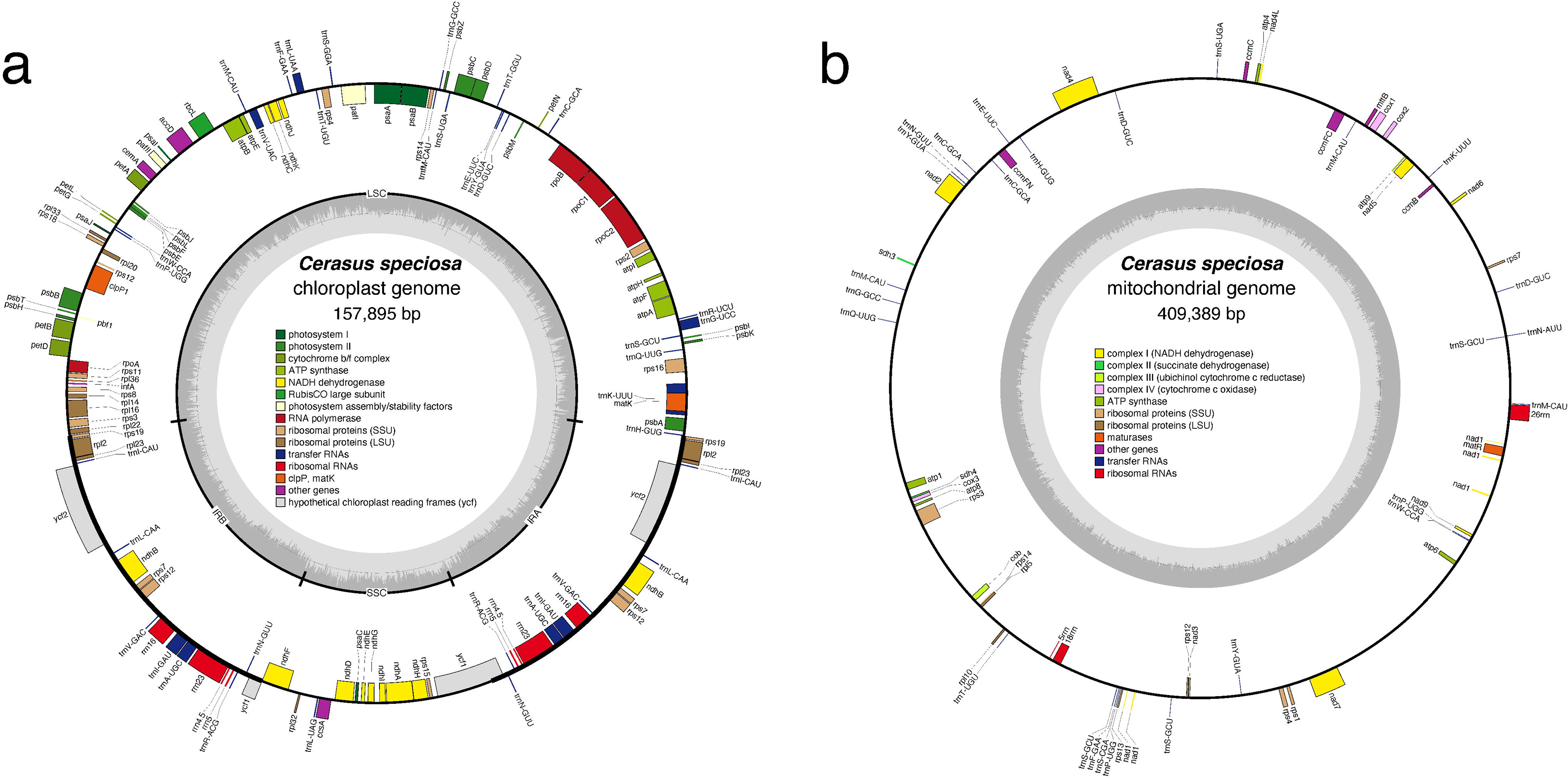
Schematic of gene annotations for the chloroplast and mitochondrial genomes of *C. speciosa*. The diagram illustrates the positional relationships between genes within the circular genomes of the chloroplasts (a) and mitochondria (b). The gene types displayed in each genome are specified in the legends corresponding to each genome. For the chloroplast genome, the track within the inner circle indicates the guanine (GC) ratio (dark color). The small single-copy (SSC), large single-copy (LSC), and inverted repeat regions (IRA and IRB) are illustrated in the inner circle.

### Mitochondrial genome assembly and annotation

Given the considerable length and abundance of repetitive sequences in plant mitochondria, which makes their assembly notably challenging compared with that of animal mitochondria, we adopted a methodology of *de novo* assembly based on PacBio long reads, with sequencing errors corrected using Illumina short reads. Initially, the subreads obtained using PacBio CLR were mapped to the complete mitochondrial genome of a closely related species using minimap2 v2.1 ^29,30^ with the parameter “-x map-pb”. The complete mitochondrial sequence of *C. avium* (NCBI RefSeq: NC_044768.1) was used as a reference. Subsequently, subreads accurately mapped to the reference were *de novo* assembled using Canu v2.0 ^57^, under the assumption that the complete mitochondrial length of *C. speciosa* is approximately 400 k, with the parameter set as genomeSize = 400 k. The resulting circular complete genome was then subjected to an improvement process using short reads, employing BWA-MEM v0.7.17 ^58^ with default parameters for mapping and Pilon v1.23 ^59^, as well as default parameters for error correction. Similar to the chloroplast genome, GeSeq was used for gene annotation. While the default settings of GeSeq were primarily employed for the annotation of the mitochondrial genome, the following adjustments were made: the “FASTA file to annotate” was set to circular, the “sequence source” was changed to “mitochondrial,” and complete mitochondrial genome of *C.* ×*yedoensis* ‘Somei-yoshino’ (NCBI RefSeq: NC_065235.1), *Cerasus* ×*kanzakura* (formerly named *Prunus kanzakura*) (NCBI RefSeq: NC_065230.1), *C. avium* (NCBI RefSeq: NC_044768.1), and *Arabidopsis thaliana* (NCBI RefSeq: NC_037304.1) were employed as the reference sequences for BLAT analysis. The final results of gene annotation were visualized as vector files in a manner similar to the complete chloroplast genome using the OGDRAW web service (Fig. 3b).

### Detecting telomeric and centromeric regions

The repetitive sequence units in the telomeric regions of *C. speciosa* genome were detected using the Telomere Identification Toolkit (tidk, https://github.com/tolkit/telomeric-identifier) with the following parameters: “tidk explore --minimum 5 --maximum 12.” This approach assumes that the repetitive sequence units in telomeric regions range from 5 to 12 bp in length. A genome-wide search conducted with “tidk search,” using the simple telomeric repeat units identified by “tidk explore” as queries, revealed telomeric repeat regions positioned at the ends in 12 out of 16 identified regions (chromosomes 1, 2, 3, and 8 exhibited telomeric repeats at both ends, whereas chromosomes 4, 5, 6, and 7 demonstrated such repeats at one end only), characterized by the sequence “CCCTAAA/TTTAGGG.” The same telomeric repeat has been reported in the closely related species *Cerasus campanulata* (formerly named *Prunus campanulata*) ^60^. These regions were identified as telomeres (Fig. 2).

To detect centromeric regions, we conducted a comprehensive scan of tandem repeat sequences of 30–1000 bp across the entire genome using the Tandem Repeats Finder (TRF) v4.09.1 ^61^ with the parameters set to “2 7 7 80 10 50 1000 -f -d -m.” The output data files generated by TRF were converted into gff files using the TRF2GFF v0.0.5. dev2+gbfe80a6 (https://github.com/Adamtaranto/TRF2GFF) software. We counted the number of repeats per period of the tandem repeat sequences detected by TRF and constructed histograms (Fig. 4a, b). In the *C. speciosa* genome, we found that repeats of 167 bp were the most common in the genome, with a total of 65,746.4 repetitions (copies ≥ 2), accounting for approximately 4.08% of the genome. Subsequently, repeats of 166 (3.03%), 332 (2.72%), 498 (2.71%), and 334 (2.34%) bp sequences were observed at high frequencies. Visualization of these detected repetitive sequences using the Integrative Genomics Viewer (IGV) v2.17.0, suggested that the minimal repeat units comprising the centromeric repetitive sequences were either 166 or 167 bp in length. Furthermore, these units appeared to have distinct distributions across different chromosomes (Fig. 4c).

**Figure 4.**
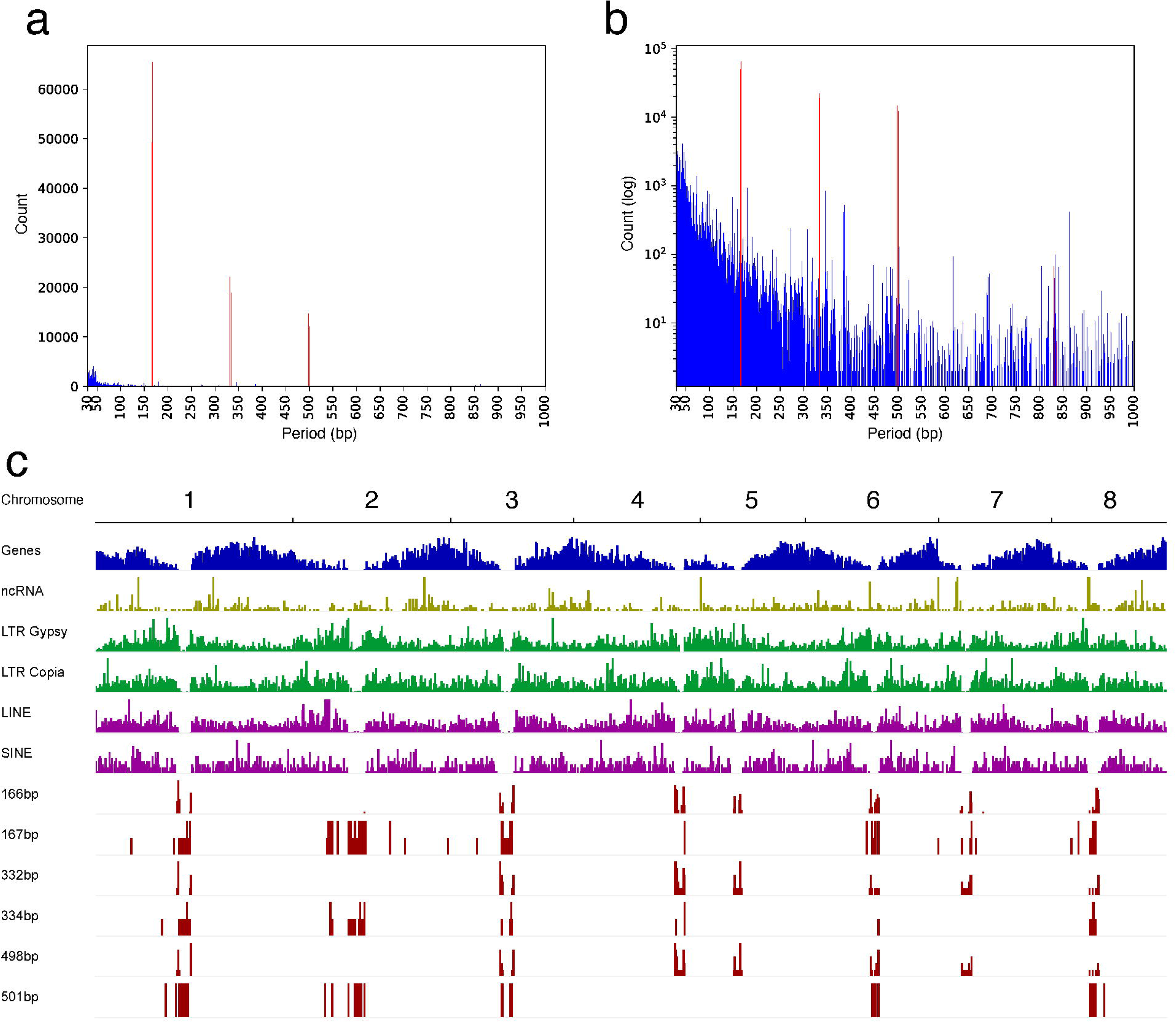
Identification of the centromeric regions via repetitive sequence analysis. (a) Frequency distribution of repetitive sequences across the genome binned by sequence length (period) in base pairs (bp). The horizontal axis represents the period of repetitive sequences, and the vertical axis represents the number of occurrences in each period. (b) Logarithmic scale representation of data shown in (a). The horizontal axis represents the period of repetitive sequences in bp, and the vertical axis represents the log-transformed count of occurrences for each period. (c) Distribution of genes, non-coding RNAs (ncRNA), LTR elements (Gypsy and Copia), LINE, SINE, and specific lengths of repetitive sequences considered to constitute centromeric structures. The horizontal axis represents the chromosomal position, and the vertical axis indicates the presence of these elements. Notably, repetitive sequences associated with centromeric structures lacked gene-rich regions, indicating putative centromeric regions.

To further investigate the detected centromeric repetitive sequences, we used the most frequently connected repetitive sequence unit (166 or 167 bp) identified in each chromosome as a monomer sequence input. We conducted a structural analysis of the regions considered as centromeres using StainedGlass v0.5 ^62^ and HiCAT v1.1.0 ^63^ (Fig. 5). The analyzed regions were specified based on the centromeric regions identified by the TRF results. For the StainedGlass analysis, a window size of 2000 bp was set. The results from StainedGlass and HiCAT revealed varying structural characteristics of the centromeric regions across chromosomes. These findings are similar to those previously reported for *Arabidopsis* ^64^.

**Figure 5.**
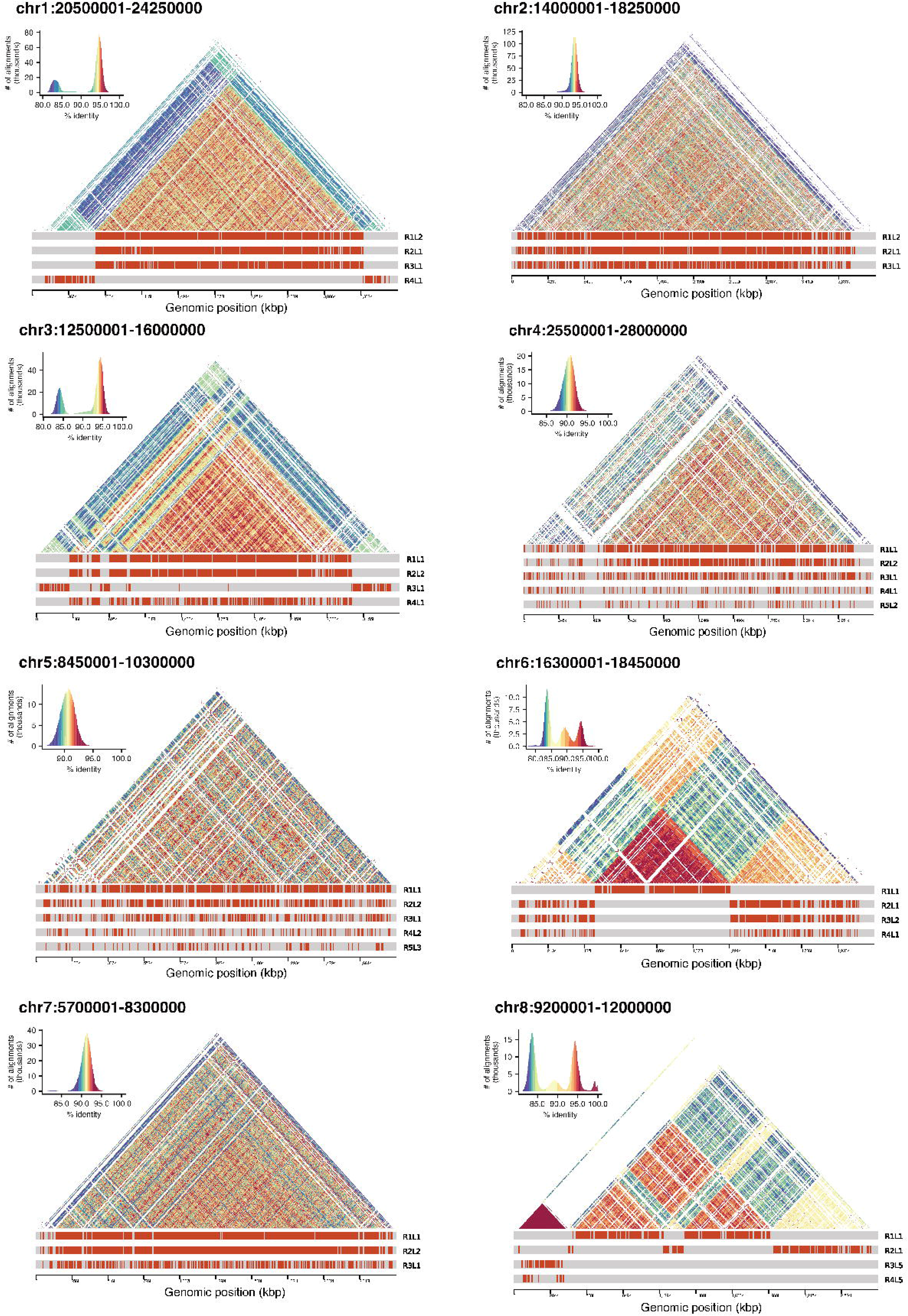
Structure and annotation of centromeric regions across chromosomes. The upper triangular sections of each chromosome represent the sequence identity of tandem repeat structures within the centromeric regions and are visualized as a heatmap. Higher similarity is indicated by warmer colors ^62^. At the bottom of the triangular sections of each chromosome, the hierarchical annotations of the estimated higher-order repeats (HORs) within the centromeric structure are displayed, showing up to the top five ranks (excluding repeats of fewer than 50 repeats) ^63^. The x-axis denotes the genomic positions in kilobase pairs (kbp), illustrating the precise location and organization of the centromeric sequences along each chromosome.

### Comparative genomic analysis

We collected the reference genome and protein sequences of *P. persica* (NCBI RefSeq GCF_000346465.2), *P. mume* (NCBI RefSeq GCF_000346735.1), *C. avium* (NCBI GenBank GCA_013416215.1), *C. campanulata* (NCBI GenBank GCA_034190965.1), and two haplotypes of *C.* ×*yedoensis* ‘Somei-yoshino’ (NCBI GenBank GCA_005406145.1) from public databases for comparative genomic analysis. Among these, the *P. mume* sequence required chromosome sorting because of a mismatch in chromosome numbers compared to other related species. For genome comparison, we constructed dot plots using D-GENIES v1.5.0 ^65^ to compare each genome with *C. speciosa*, and aligned the sequences using Minimap2 v2.24 ^30^ implemented in D-GENIES, with the “"Few repeats” option specified (Fig. 6). The genomes of related species lack regions presumed to be centromeres, resulting in observable gaps when compared to our genome. Structural differences among the compared species were generally minimal, with *C. campanulata* showing a remarkably similar genomic structure, suggesting a close phylogenetic relationship. In the comparisons with *C.* ×*yedoensis* ‘Somei-yoshino’, some regions exhibited inversions and translocations, but these findings are likely artifacts due to the low quality of the pseudomolecule assembly of this genome. Furthermore, we performed an interspecies synteny analysis using the protein sequences of *P. persica* and *C. campanulata,* which are particularly well-assembled among closely related species. For the detection of synteny regions, we employed MCscanX ^66^, preceded by a homology search of the protein sequences from each genome using Protein-Protein BLAST v2.12.0+, with an e-value cutoff of “1e-10.” The results from MCscanX were visualized using SynVisio (https://synvisio.github.io) (Fig. 7).

**Figure 6.**
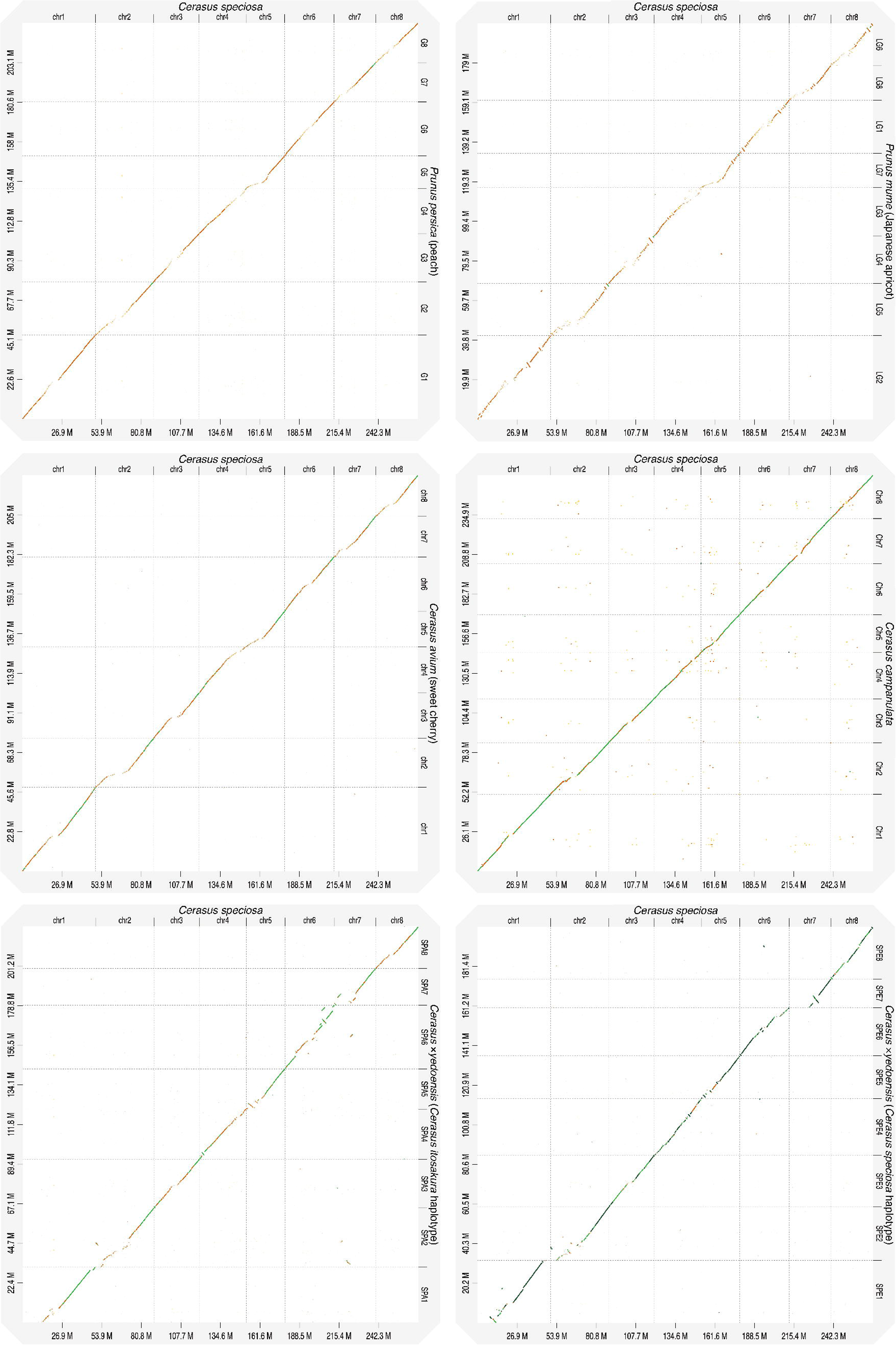
Synteny analysis of *C. speciosa* and other reference genomes. The dot plot illustrates the syntenic relationship between the eight chromosomes of *C. speciosa* (x-axis) and the contigs of the reference genome (y-axis). Each dot represents a conserved genomic block between two genomes. The diagonal green line indicates collinear regions, suggesting high conservation and alignment among the compared genomes.

**Figure 7.**
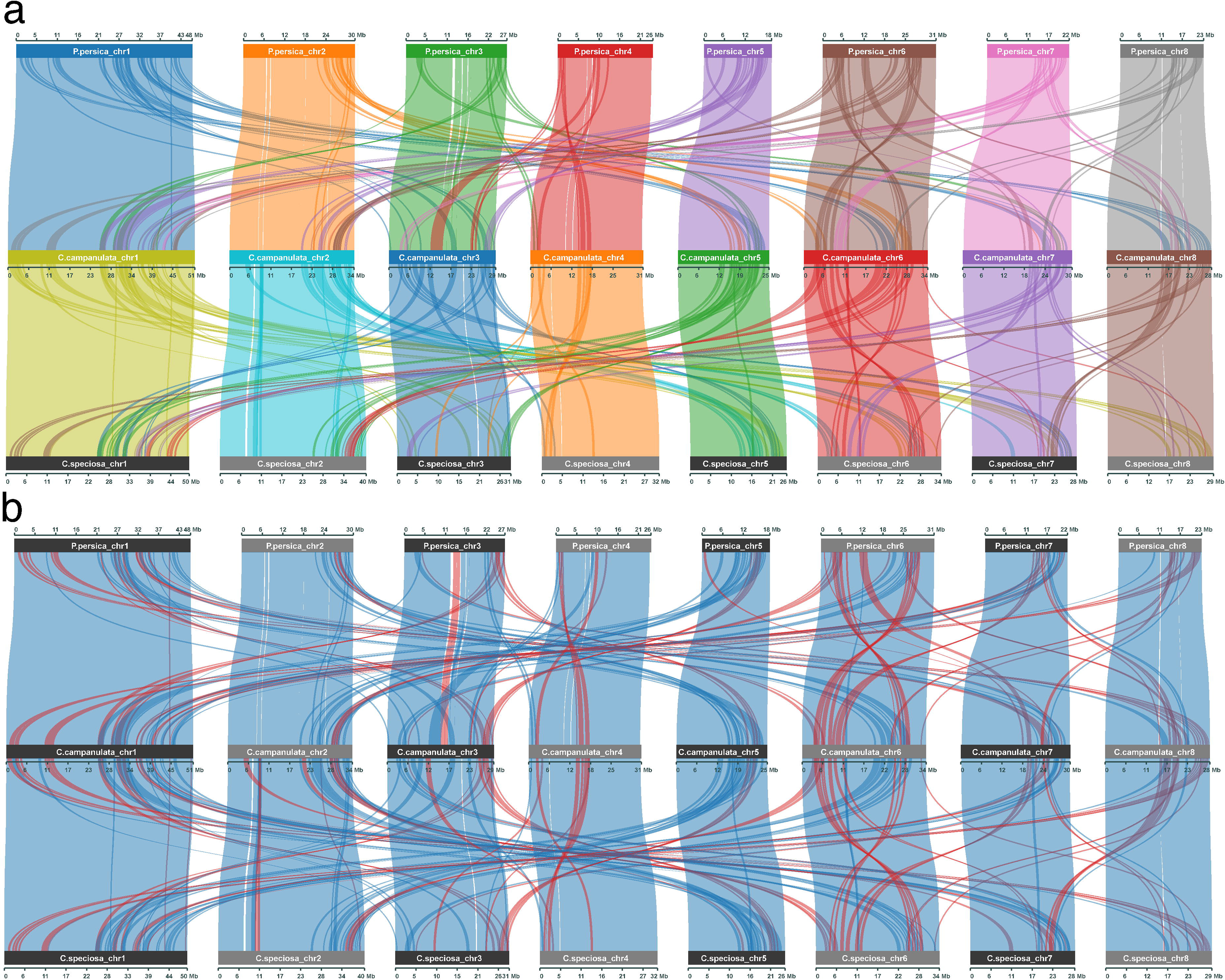
Comparative analysis of protein-coding genes in *P. persica*, *C. campanulata,* and *C. speciosa,* and chromosomal distributions of protein-coding genes in *P. persica*, *C. campanulata*, and *C. speciosa*. In the upper panel (a), each color represents a different chromosome, illustrating the collinearity patterns of genes across different chromosomes. In the lower panel (b), each line shows the orientation of the collinearity patterns of the genes. The figure shows whether the collinearity patterns of the genes were in forward or reverse orientation. Genes in red indicate a reverse orientation.

## Data Records

All raw data used in this study for genome assembly (Illumina short reads, PacBio CLR reads, PacBio HiFi reads, and Hi-C reads) were deposited in the DNA Data Bank of Japan (DDBJ) under BioProject accession number PRJDB17512. As DDBJ is a member of the International Nucleotide Sequence Database Collaboration, the data are accessible via the National Center for Biotechnology Information (NCBI) and European Nucleotide Archive (ENA). Accession numbers for the data are as follows: DRR532494 for Illumina short reads, DRR532493 for PacBio CLR reads, DRR529729 for PacBio HiFi reads, and DRR529730 for Hi-C reads. The RNA-seq data from the Illumina short reads used for annotating the assembled genome were also registered under the same BioProject accession number PRJDB17512, with individual data for each tissue type under accession numbers DRR530280–DRR530285. The assembled genome sequences were deposited in the DDBJ under the accession numbers AP029589–AP029596 for each chromosome. Additionally, the chloroplast and mitochondrial genomes were registered under the accession numbers AP029597 and AP029598, respectively. The assembled genome sequences and gene annotation data were submitted to the Genome Database for Rosaceae (GDR; www.rosaceae.org) under analysis URL (https://www.rosaceae.org/Analysis/20425168). These datasets are publicly available from the cherry genome database (https://sakura.nig.ac.jp/genome/). The monomer sequence of the centromeric region used for the centromeric region structure and hierarchy annotation analyses is available on Figshare with the doi [10.6084/m9.figshare.26046853].

## Technical Validation

The quality of the *C. speciosa* genome assembly was assessed using four methodologies. The first method employed was an evaluation of completeness using the BUSCO assessment with the “eudicots_odb10” lineage dataset (n = 2326), representative of eudicots. The final BUSCO assessment revealed a completeness rate of 98.4% (n = 2289), comprising 95.9% (n = 2231) single-copy, 2.5% (n = 58) duplicated, 0.5% (n = 11) fragmented, and 1.1% (n = 26) missing BUSCOs. Additionally, a second method involves comfort evaluation, which is proposed to be faster and more accurate than BUSCO. Utilizing the same “eudicots_odb10” BUSCO gene set, evaluation of compleasm showed a completeness of 99.1% (n = 2306), with 97.0% (n = 2257) single-copy, 2.1% (n = 49) duplicated, 0.4% (n = 9) fragmented, and 0.5% (n = 11) missing BUSCOs. Although the evaluation using Compleasm indicated a higher quality than that of BUSCO, the BUSCO assessment itself sufficiently demonstrated the high quality of the genome sequence produced in this study. Third, we used Inspector to assess our novel genome sequences by generating alignments of PacBio HiFi sequencing reads, which were used for genome assembly, with the assembled genome sequences. The mapping rates from reads-to-contigs and QV were 99.99% and 35.89 (corresponding to 99.97% accuracy), respectively. Finally, we employed Merqury, a reference-free, *k*-mer-based genome assembly evaluation approach, for validation. This method involves measuring the *k*-mer spectrum from a read set used for assembly, thereby enabling independent evaluation without the use of a reference. To construct the *k*-mer database, we specified *k*=21 using PacBio HiFi reads from Meryl. The results from this approach further indicated that our assembly data exhibited high-quality metrics (QV > 67 with an error rate of 1.80 × 10^-7^). A comprehensive assessment using all these methodologies demonstrated the completeness of the *C. speciosa* genome assembly (Table 2).

## Code availability

No specific code or script was used in the study. All the software used in this study were executed according to their manuals and protocols. Details of the versions and parameters used in this study are described in the Methods section, with any unspecified parameters set to the default for the respective versions.

## Acknowledgements

We thank Motoko Nihei and Kumiko Masuyama for their technical assistance in this study. We express our sincere gratitude to Dr. Tetsuya Hori and Dr. Tatsuo Fukagawa for their invaluable suggestions regarding centromeric region analysis. We also express our sincere gratitude to Oshima town, the Tokyo Metropolitan Government, and the Agency for Cultural Affairs for their cooperation in the sampling of the special natural monument “Sakurakkabu.” This work was supported by grants from (Research Organization of Information and Systems). We would like to thank Editage (www.editage.jp) for English language editing.

## Author contributions

T. Koide conceived the project; K.F., B.B.B., T. Kishida, and M.T. analyzed the data; A.T. performed the sequencing; T. Koide, K.F., and T. Katsuki performed the sampling; T. Koide, A.T., and Y.S. extracted the genomic DNA and RNA. N.K. and S.K. created cherry genome databases. A.K., K.N., H.N., H.Y., K.U., and T. O. C. S. were involved in the planning and key discussions throughout this project. The initial draft of the manuscript was authored by K.F., who incorporated detailed feedback from all the authors. All authors approved the final version of the manuscript.

Sakura 100 Genome Consortium (full list): Kazumichi Fujiwara^1^, Atsushi Toyoda^2^, Bhim B. Biswa^1,3^, Takushi Kishida^4^, Momi Tsuruta^5^, Yasukazu Nakamura^3,6^, Noriko Kimura^7^, Shoko Kawamoto^3,7^, Yutaka Sato^3,8^, Toshio Katsuki^9^, Tsuyoshi Koide^1,3^, Akatsuki Kimura^10^, Ken-Ichi Nonomura^11^, Hironori Niki^12^, Hiroyuki Yano^13^, Kinji Umehara^14^, Tazro Ohta^15^, Chikahiko Suzuki^16^.

^10^Cell Architecture Laboratory, National Institute of Genetics, Mishima, Japan; ^11^Plant Cytogenetics Laboratory, National Institute of Genetics, Mishima, Japan; ^12^Microbial Physiology Laboratory, National Institute of Genetics, Mishima, Japan; ^13^Invertebrate Genetics Laboratory, National Institute of Genetics, Mishima, Japan; ^14^Association for Propagation of the Knowledge of Genetics, Mishima, Japan; ^15^Institute for Advanced Academic Research, Chiba University, Chiba, Japan; ^16^Culture and Information, Gunma Prefectural Women’s University, Tamamuramachi, Japan.

## Competing interests

The authors declare no competing interests.

## Notes

### Competing Interest Statement

The authors have declared no competing interest.

## Reference

1 Ohba, H. in Flora of Japan Vol. IIb (eds Kunio Iwatsuki, Takasi Yamazaki, David E. Boufford, & Hideaki Ohba) 128–144 (Kodansha, 2001).

2 Katsuki, T. & Iketani, H. Nomenclature of Tokyo cherry (*Cerasus* × *yedoensis* ‘Somei yoshino’, Rosaceae) and allied interspecific hybrids based on recent advances in population genetics. Taxon 65, 1415–1419 (2016).

3 Arakawa, H. Twelve centuries of blooming dates of the cherry blossoms at the city of Kyoto and its own vicinity. Geofisica Pura e Applicata 30, 147–150 (1955).

4 Kuitert, W. & Peterse, A. Japanese flowering cherries. (Timber Press, 1999).

5 Aono, Y. & Kazui, K. Phenological data series of cherry tree flowering in Kyoto, Japan, and its application to reconstruction of springtime temperatures since the 9th century. International Journal of Climatology 28, 905-914 (2007).

6 Kawasaki, T. Flowering cherries of Japan. (Yama-Kei Publisher, 1993). (in Japanese)

7 Ikeda, H., Iketani, H. & Katsuki, T. in Wild Flowers of Japan, revised new edition Vol. 3 (eds Hiroyoshi Ohashi et al.) 23–88 (Heibonsha, 2017). (in Japanese)

8 Katsuki, T. A New Species, *Cerasus kumanoensis* from the Southern Kii Peninsula, Japan. Acta Phytotaxonomica et Geobotanica 69, 119–126 (2018).

9 Miyoshi, M. Japanische Bergkirschen, ihre Wildformen und Kulturrassen. Journal College of Science Imperial University of Tokyo 34, 175 (1916). (in German)

10 Wilson, E. H. The cherries of Japan. (University press, 1916).

11 Kato, S. et al. Clone identification in Japanese flowering cherry (*Prunus* subgenus *Cerasus*) cultivars using nuclear SSR markers. Breed Sci 62, 248–255 (2012).

12 Kato, S. et al. Origins of Japanese flowering cherry (*Prunus* subgenus *Cerasus*) cultivars revealed using nuclear SSR markers. Tree Genetics & Genomes 10, 477–487 (2014).

13 Takenaka, Y. The origin of the yoshino cherry tree. The Journal of Heredity 54, 207–211 (1963).

14 Innan, H., Terauchi, R., Miyashita, N. T. & Tsunewaki, K. DNA fingerprinting study on the intraspecific variation and the origin of *Prunus yedoensis* (Someiyoshino). The Japanese Journal of Genetics 70, 185–196 (1995).

15 Shirasawa, K. et al. Phased genome sequence of an interspecific hybrid flowering cherry, ‘Somei-Yoshino’ (*Cerasus* × *yedoensis*). DNA Res 26, 379–389 (2019).

16 Takaishi, K. Studies on the coumarin components in the leaves of cherry-tree. Yakugaku Zasshi 88, 1467–1471 (1968).

17 Ohba, H., Kawasaki, T., Kihara, H. & Tanaka, H. Flowering cherries of Japan. New Edition edn, (Yama-Kei Publishers, 2007). (in Japanese)

18 Oginuma, K. & Tanaka, R. Karyomorphological studies on some cherry trees in Japan. The Journal of Japanese Botany 51, 104–109 (1976).

19 Miyoshi, M. Botanical Natural Monuments of Tokyo and Other Prefectures. 1–6 (The Ministry of Education, Science, Sports and Culture, Japan, 1936). (in Japanese)

20 Carlson, J. E. et al. Segregation of random amplified DNA markers in F1 progeny of conifers. Theoretical and Applied Genetics 83, 194–200 (1991).

21 Baid, G. et al. DeepConsensus improves the accuracy of sequences with a gap-aware sequence transformer. Nat Biotechnol 41, 232–238 (2023).

22 Chen, S., Zhou, Y., Chen, Y. & Gu, J. fastp: an ultra-fast all-in-one FASTQ preprocessor. Bioinformatics 34, i884–i890 (2018).

23 Kokot, M., Dlugosz, M. & Deorowicz, S. KMC 3: counting and manipulating k-mer statistics. Bioinformatics 33, 2759–2761 (2017).

24 Vurture, G. W. et al. GenomeScope: fast reference-free genome profiling from short reads. Bioinformatics 33, 2202–2204 (2017).

25 Ranallo-Benavidez, T. R., Jaron, K. S. & Schatz, M. C. GenomeScope 2.0 and Smudgeplot for reference-free profiling of polyploid genomes. Nat Commun 11, 1432 (2020).

26 Cheng, H., Concepcion, G. T., Feng, X., Zhang, H. & Li, H. Haplotype-resolved de novo assembly using phased assembly graphs with hifiasm. Nat Methods 18, 170–175 (2021).

27 Cheng, H. et al. Haplotype-resolved assembly of diploid genomes without parental data. Nat Biotechnol 40, 1332–1335 (2022).

28 Guan, D. et al. Identifying and removing haplotypic duplication in primary genome assemblies. Bioinformatics 36, 2896–2898 (2020).

29 Li, H. New strategies to improve minimap2 alignment accuracy. Bioinformatics 37, 4572–4574 (2021).

30 Li, H. Minimap2: pairwise alignment for nucleotide sequences. Bioinformatics 34, 3094–3100 (2018).

31 Altschul, S. F., Gish, W., Miller, W., Myers, E. W. & Lipman, D. J. Basic local alignment search tool. Journal of Molecular Biology 215, 403–410 (1990).

32 Manni, M., Berkeley, M. R., Seppey, M., Simao, F. A. & Zdobnov, E. M. BUSCO Update: Novel and Streamlined Workflows along with Broader and Deeper Phylogenetic Coverage for Scoring of Eukaryotic, Prokaryotic, and Viral Genomes. Mol Biol Evol 38, 4647–4654 (2021).

33 Kriventseva, E. V. et al. OrthoDB v10: sampling the diversity of animal, plant, fungal, protist, bacterial and viral genomes for evolutionary and functional annotations of orthologs. Nucleic Acids Res 47, D807–D811 (2019).

34 Zdobnov, E. M. et al. OrthoDB in 2020: evolutionary and functional annotations of orthologs. Nucleic Acids Res 49, D389–D393 (2021).

35 Huang, N. & Li, H. compleasm: a faster and more accurate reimplementation of BUSCO. Bioinformatics 39 (2023).

36 Chen, Y., Zhang, Y., Wang, A. Y., Gao, M. & Chong, Z. Accurate long-read de novo assembly evaluation with Inspector. Genome Biol 22, 312 (2021).

37 Ou, S. et al. Benchmarking transposable element annotation methods for creation of a streamlined, comprehensive pipeline. Genome Biol 20, 275 (2019).

38 Contreras-Moreira, B. et al. K-mer counting and curated libraries drive efficient annotation of repeats in plant genomes. Plant Genome 14, e20143 (2021).

39 Nussbaumer, T. et al. MIPS PlantsDB: a database framework for comparative plant genome research. Nucleic Acids Res 41, D1144–1151 (2013).

40 Amselem, J. et al. RepetDB: a unified resource for transposable element references. Mob DNA 10, 6 (2019).

41 Wicker, T., Matthews, D. E. & Keller, B. TREP:a database for Triticeae repetitive elements. Trends in Plant Science 7, 561–562 (2002).

42 Chan, P. P., Lin, B. Y., Mak, A. J. & Lowe, T. M. tRNAscan-SE 2.0: improved detection and functional classification of transfer RNA genes. Nucleic Acids Res 49, 9077–9096 (2021).

43 Lagesen, K. et al. RNAmmer: consistent and rapid annotation of ribosomal RNA genes. Nucleic Acids Res 35, 3100–3108 (2007).

44 Kalvari, I. et al. Rfam 14: expanded coverage of metagenomic, viral and microRNA families. Nucleic Acids Res 49, D192–D200 (2021).

45 Kozomara, A., Birgaoanu, M. & Griffiths-Jones, S. miRBase: from microRNA sequences to function. Nucleic Acids Res 47, D155–D162 (2019).

46 Flynn, J. M. et al. RepeatModeler2 for automated genomic discovery of transposable element families. Proc Natl Acad Sci U S A 117, 9451–9457 (2020).

47 Gabriel, L., et al. BRAKER3: Fully automated genome annotation using RNA-seq and protein evidence with GeneMark-ETP, AUGUSTUS and TSEBRA. bioRxiv (2024).

48 Bruna, T., Lomsadze, A. & Borodovsky, M. A new gene finding tool GeneMark-ETP significantly improves the accuracy of automatic annotation of large eukaryotic genomes. bioRxiv (2024).

49 Grabherr, M. G. et al. Full-length transcriptome assembly from RNA-Seq data without a reference genome. Nat Biotechnol 29, 644–652 (2011).

50 Kim, D., Paggi, J. M., Park, C., Bennett, C. & Salzberg, S. L. Graph-based genome alignment and genotyping with HISAT2 and HISAT-genotype. Nat Biotechnol 37, 907–915 (2019).

51 Stanke, M., Diekhans, M., Baertsch, R. & Haussler, D. Using native and syntenically mapped cDNA alignments to improve de novo gene finding. Bioinformatics 24, 637–644 (2008).

52 Hart, A. J. et al. EnTAP: Bringing faster and smarter functional annotation to non-model eukaryotic transcriptomes. Mol Ecol Resour 20, 591–604 (2020).

53 Buchfink, B., Xie, C. & Huson, D. H. Fast and sensitive protein alignment using DIAMOND. Nat Methods 12, 59–60 (2015).

54 Jin, J. J. et al. GetOrganelle: a fast and versatile toolkit for accurate de novo assembly of organelle genomes. Genome Biol 21, 241 (2020).

55 Tillich, M. et al. GeSeq - versatile and accurate annotation of organelle genomes. Nucleic Acids Res 45, W6–W11 (2017).

56 Greiner, S., Lehwark, P. & Bock, R. OrganellarGenomeDRAW (OGDRAW) version 1.3.1: expanded toolkit for the graphical visualization of organellar genomes. Nucleic Acids Res 47, W59-W64 (2019).

57 Koren, S. et al. Canu: scalable and accurate long-read assembly via adaptive k-mer weighting and repeat separation. Genome Res 27, 722–736 (2017).

58 Li, H. & Durbin, R. Fast and accurate short read alignment with Burrows-Wheeler transform. Bioinformatics 25, 1754–1760 (2009).

59 Walker, B. J. et al. Pilon: an integrated tool for comprehensive microbial variant detection and genome assembly improvement. PLoS One 9, e112963 (2014).

60 Hu, Y., Feng, C., Wu, B. & Kang, M. A chromosome-scale assembly of the early-flowering *Prunus campanulata* and comparative genomics of cherries. Sci Data 10, 920 (2023).

61 Benson, G. Tandem repeats finder: a program to analyze DNA sequences. Nucleic Acids Res 27, 573–580 (1999).

62 Vollger, M. R., Kerpedjiev, P., Phillippy, A. M. & Eichler, E. E. StainedGlass: interactive visualization of massive tandem repeat structures with identity heatmaps. Bioinformatics 38, 2049–2051 (2022).

63 Gao, S. et al. HiCAT: a tool for automatic annotation of centromere structure. Genome Biol 24, 58 (2023).

64 Naish, M. et al. The genetic and epigenetic landscape of the Arabidopsis centromeres. Science 374, eabi7489 (2021).

65 Cabanettes, F. & Klopp, C. D-GENIES: dot plot large genomes in an interactive, efficient and simple way. PeerJ 6, e4958 (2018).

66 Wang, Y. et al. MCScanX: a toolkit for detection and evolutionary analysis of gene synteny and collinearity. Nucleic Acids Res 40, e49 (2012).

